# A high-throughput method for measuring critical thermal limits of leaves by chlorophyll imaging fluorescence

**DOI:** 10.1101/2020.09.21.306886

**Authors:** Pieter A. Arnold, Verónica F. Briceño, Kelli M. Gowland, Alexandra A. Catling, León A. Bravo, Adrienne B. Nicotra

## Abstract

Plant thermal tolerance is a crucial research area as the climate warms and extreme weather events become more frequent. Leaves exposed to temperature extremes have inhibited photosynthesis and will accumulate damage to photosystem II (PSII) if tolerance thresholds are exceeded. Temperature-dependent changes in basal chlorophyll fluorescence (*T-F*_0_) can be used to identify the critical temperature at which PSII is inhibited. We developed and tested a high-throughput method for measuring the critical temperatures for PSII at low (*CT*_MIN_) and high (*CT*_MAX_) temperatures using a Maxi-Imaging fluorimeter and a thermoelectric Peltier plate heating/cooling system. We examined how experimental conditions: wet *vs* dry surfaces for leaves and heating/cooling rate, affect *CT*_MIN_ and *CT*_MAX_ across four species. *CT*_MAX_ estimates were not different whether measured on wet or dry surfaces, but leaves were apparently less cold tolerant when on wet surfaces. Heating/cooling rate had a strong effect on both *CT*_MAX_ and *CT*_MIN_ that was species-specific. We discuss potential mechanisms for these results and recommend settings for researchers to use when measuring *T-F*_0_. The approach that we demonstrated here allows the high-throughput measurement of a valuable ecophysiological parameter that estimates the critical temperature thresholds of leaf photosynthetic performance in response to thermal extremes.

## Introduction

Understanding both vulnerability and tolerance limits of plants to thermal extremes is a priority for plant biology research as the Earth’s climate continues to change, thereby exposing these sessile organisms to increased thermal stress (O’Sullivan *et al*. 2017; IPCC 2018; Geange *et al*. 2021). Thermal stress disrupts and inhibits physiological processes (Goraya *et al*. 2017), induces protective and repair mechanisms (Sung *et al*. 2003; Goh *et al*. 2012), leads to declines in plant performance, and threatens survival (Zinn *et al*. 2010; Bita and Gerats 2013). Plant photosynthesis is sensitive to thermal stress and has distinct limits beyond which photosynthetic assimilation is inhibited and tissue damage can occur (e.g., Neuner and Pramsohler 2006; Sukhov *et al*. 2017). The temperature sensitivity of photosynthesis is in part derived from the thermally-dependent stability of protein-pigment complexes in the light harvesting complex II (LHCII) of photosystem II (PSII) of the thylakoid membrane in chloroplasts (Ilík *et al*. 2003), which are integral to the photosynthetic electron transport chain (Berry and Björkman 1980; Allakhverdiev *et al*. 2008; Mathur *et al*. 2014).

Chlorophyll fluorimetry has become a widely used tool for assessing the thermal limits of photosynthesis for both cold and heat tolerance (Geange *et al*. 2021). Chlorophyll can dissipate absorbed light energy via photochemistry or re-emit it as heat energy or fluorescence (Baker 2008; Murchie and Lawson 2013). A dark-adapted leaf exposed to a low-intensity modulated measuring light, which does not induce electron transport, emits a minimal amount of chlorophyll-*a* fluorescence from LHCII, called *F*_0_ (Yamane *et al*. 1997). Under more intense or actinic light, processes that are highly dynamic and sensitive to other factors but not well correlated with the viability of the photosynthetic tissue cannot be isolated from the measurement of the temperature dependence (thermal stability) of chlorophyll fluorescence (Schreiber *et al*. 1995; Logan *et al*. 2007). To assess the thermal stability limits of LHCII, plant ecophysiologists typically measure the temperature-dependent change in basal chlorophyll-*a* fluorescence (*T-F*_0_) to determine the critical temperature threshold (*T*_crit_), denoted by a sudden increased in *F*_0_ at which PSII begins to inactivate (e.g., Schreiber and Berry 1977; Berry and Björkman 1980; Briantais *et al*. 1996; Knight and Ackerly 2002; Ilík *et al*. 2003; Hüve *et al*. 2006; Neuner and Pramsohler 2006; O’Sullivan *et al*. 2013; O’Sullivan *et al*. 2017; Zhu *et al*. 2018). *F*_0_ is a fluorescence parameter that can be measured rapidly and continuously throughout heating or cooling in darkness, without the need of a saturating pulse and re-dark adaptation as for *F*_V_/*F*_M_ measurements that are commonly used to detect photosynthetic inhibition.

One critique of *T-F*_0_ measurements and *T*_crit_ determination is that they are conducted on detached leaves. Detaching leaves to expose them to a precisely controlled and measured thermal surface is usually, but not always, a necessary component of this trait measurement. While modern chlorophyll fluorescence imaging systems can be used on attached leaves, simultaneously heating or cooling these leaves precisely while measuring multiple leaf samples remains logistically complex, especially for ecological applications. Leaf detachment can affect leaf hydration and fluorescence through reduced PSII activity, ionic leakage, and oxidations compared to attached leaves (Potvin 1985; Smillie *et al*. 1987). Leaf dehydration could be problematic for certain species if leaves are sampled long before they are assessed for *T*_crit_ or if they are measured as leaf sections or discs. To avoid dehydration during the *T-F*_0_ measurement, a wet surface, such as damp paper surface as in Knight and Ackerly (2002), could physically impair evaporation by saturating the atmosphere surrounding the leaf. However, it is not clear whether a wet surface interferes with the *T-F*_0_ measurement or how it might affect the *T*_crit_ value compared to using a dry surface.

A great advantage of using temperature-dependent changes in chlorophyll fluorescence and a thermoelectric plate is that both cold and heat tolerance limits of leaves can be measured with much of the same equipment. However, the protocol may need to be altered slightly because cold transitions in nature occur much more slowly than heat transitions, which may induce different mechanisms in response to thermal stress. For example, leaf temperature can rapidly increase during a lull in wind speed, far exceeding ambient temperature on a hot and sunny day (Vogel 2009; Leigh *et al*. 2012). On a cold frosty night, even considering air temperature stratification, the rate of leaf temperature cooling rarely exceeds 5°C h^-1^, especially below freezing (Sakai and Larcher 1987). Therefore, the ‘standard’ protocols for measuring *T*_crit_ typically change temperature much faster for heat tolerance than for cold tolerance. While this approach is justified by rates observed in natural systems, the first published application of the *T-F*_0_ technique (Schreiber and Berry 1977) used an apparently arbitrary ‘slow’ heating rate of 1°C min^-1^ (i.e., 60°C h^-1^). Subsequently, while many studies followed suit, a vast range of heating/cooling rates have been applied (see Table S1, available as Supplementary Material to this paper), often with little justification. We have known for decades that different rates of heating and cooling can affect the *T-F*_0_ curve and shift the *T*_crit_ value by at least 2°C (Bilger *et al*. 1984; Frolec *et al*. 2008). Therefore, studies employing *T-F*_0_ methods for measuring thermal tolerance limits that use different heating/cooling rates might not be directly comparable, even within a given species. Further, it is reasonable to expect that plant species might exhibit different responses to variation in methodology.

Here, we present a practical, high-throughput method for measuring *T*_crit_ with a Pulse Amplitude Modulated (PAM) chlorophyll fluorescence imaging system that measures *F*_0_ in real time as a thermoelectric Peltier plate with leaf samples is heated or cooled to thermal extremes. We then investigate variations of easily controllable variables of the standard experimental protocol that could affect thermal tolerance limit estimates. We sought to determine the effects of wet *vs* dry surface and heating/cooling rate on *T*_crit_ estimates for both the heat tolerance limit (hereafter referred to as critical maximum temperature; *CT*_MAX_) and the cold tolerance limit (hereafter referred to as critical minimum temperature; *CT*_MIN_) of leaf thermal stability of species with different growth forms. By comparing among these species, we also determined whether the effects of the two experimental variables could be generalised for different growth forms of plants that originate from different conditions. In doing so, we advise researchers on what we consider to be a pragmatic approach to measuring leaf thermal tolerance using chlorophyll imaging fluorescence, at a time when improved understanding of plant tolerance to thermal extremes is needed for cultivated and wild species alike.

## Materials and Methods

### Species description and leaf samples

We chose plant species that represented diverse growth habits and leaf morphology (in surface characteristics and leaf thickness) to make simple interspecific comparisons while testing the *T-F*_0_ method. *Wahlenbergia ceracea* Lothian (Campanulaceae) waxy bluebell is a small perennial herb that is sparsely distributed across south-eastern Australia. We grew F2 generation *W. ceracea* plants under controlled glasshouse conditions (20/15°C set day/night temperatures) and leaves from mature plants were used for all experiments. Seed stock originated from Kosciuszko National Park, NSW, Australia (36.432°S, 148.338°E) that was collected in 2015 and 2016. *Melaleuca citrina* (Curtis) Dum. Cours. (Myrtaceae) common red bottlebrush were used for all experiments. This species is native to south-eastern Australia but also distributed as a cosmopolitan plant. Sampled individuals were growing as native shrubs at The Australian National University, ACT, Australia (35.279°S, 149.118°E). *Quercus phellos* L. (Fagaceae) willow oak trees were used only in the heat tolerance component of the surface wetness experiment, prior to the abscission of leaves in autumn. This deciduous species is native to North America and sampled individuals were growing as tall, shady ornamental trees at The Australian National University, ACT, Australia (35.277°S, 149.115°E). *Escallonia rubra* var. ‘pink pixie’ (Ruiz & Pav.) Pers. (Escalloniaceae) pink escallonia were used for the cold tolerance component of the surface wetness experiment and the heating/cooling rate experiment in place of *Q. phellos* after the former shed its leaves. *Escallonia rubra* is native to South America and sampled individuals were growing as dense ornamental shrubs at The Australian National University, ACT, Australia (35.277°S, 149.117°E).

All measurements were taken between February and October 2019. Due to the variation in species availability across experiments and the potential effects of seasonal change on absolute tolerance values, we consider each experiment separately and do not draw comparisons across surface wetness and heating/cooling rate experiments. Assays (surface wetness or heating/cooling rates for heat or cold tolerance assays) were conducted on replicate days to control for potential effects of day. Leaves selected for measurement were fully expanded, visually free of damage and discolouration, and within two leaf pairs of a growing stem tip on an intact and healthy stem. Although leaf age could not be determined directly, these criteria allowed us to select leaves from the same cohort and of similar condition. Leaves were excised between 0900 and 1300 hours, placed in sealed bags, and then taken to the lab in an insulated container, where they were always used for *T-F*_0_ measurements within 30 minutes of initial collection.

### Temperature-dependent change in chlorophyll fluorescence (T-F_0_) measurement

Leaf samples were attached to white filter paper (125 × 100 mm) with double-sided tape. We placed the filter paper with leaves on a Peltier plate (CP-121HT; TE-Technology, Inc., Michigan, USA; 152 × 152 mm surface) that was controlled by a bi-polar proportional-integral-derivative temperature controller (TC-36-25; TE-Technology, Inc.) and powered by a fixed-voltage power supply (PS-24-13; TE-Technology, Inc.). The Peltier plate uses four direct-contact thermoelectric modules that can both cool and heat the plate, which with a MP-3193 thermistor (TE-Technology, Inc.) the plate had potential thermal limits of –20°C and 100°C. LabVIEW-based control software (National Instruments, Texas, USA) was adapted to control heating or cooling rate using source code available from TE-Technology, Inc. based on the supplied user interface. The Peltier plate maintained a stable set temperature within ± 0.1°C (precision) and ± 1°C tolerance across the plate surface. We attached two type-T thermocouples to the underside of two randomly selected leaves on the plate as representative measures of leaf temperatures. Thermocouple temperature data were recorded every 10 s by a dual-channel data logger (EL-GFX-DTC; Lascar Electronics Ltd., Salisbury, UK) and the mean temperature of the two thermocouples was used for all leaf temperature calculations. Because the two thermocouples measured temperatures of two single leaves per experimental run, we were able to extract a small subset of ice nucleation temperatures (*NT*) using the temperature of the first exothermic reaction in cold tolerance assays. The Peltier plate assembly height was controlled by a laboratory scissor-jack to fit within an aluminium frame at an ideal height below the fluorescence camera (Fig. 1*a*). Heavy double-glazed glass was placed on top of the leaf samples on the plate to compress samples against the plate surface to ensure maximum contact and create a thermal buffer to ensure close matching of leaf and plate temperatures. In addition to greater thermal buffering relative to standard glass, double-glazed glass avoids condensation that might lead to erroneous measurements of *F*_0_. All areas of both the Peltier plate and glass that were outside of the filter paper area were blacked out with heat-resistant black electrical tape to remove ambient light reflection and interference.

**Fig 1.**
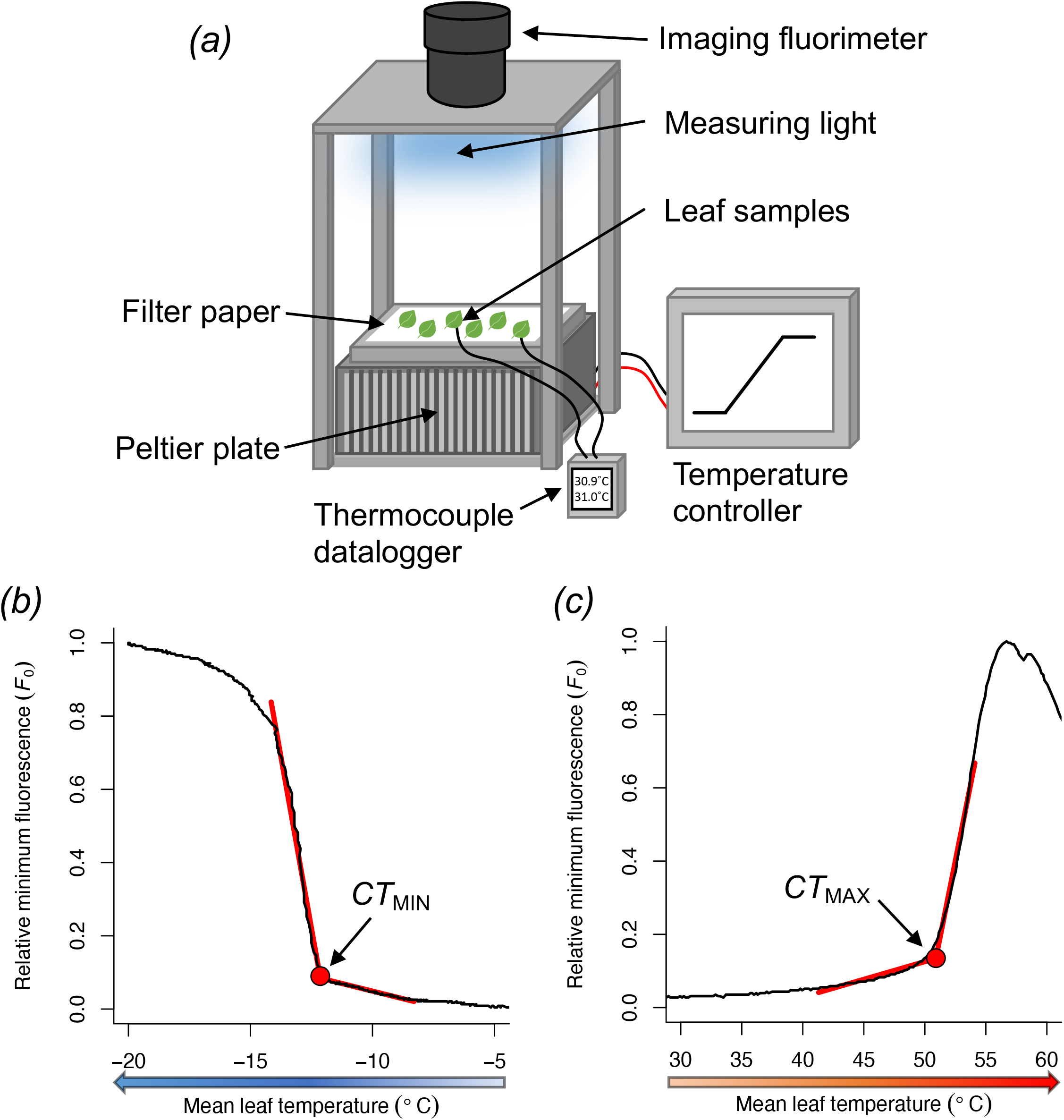
Experimental system for measuring thermal tolerance limits and representative temperature-dependent chlorophyll fluorescence curves (*T-F*_0_). (*a*) The Peltier plate-Maxi-Imaging fluorimeter setup for measuring leaf thermal tolerance limits. (*b*) Representative *T-F*_0_ curve for *CT*_MIN_ (inflection point is the *T*_crit_) where leaf sample temperature (°C) decreases to a point below freezing where the leaf rapidly emits more fluorescence (*F*_0_, relative units), indicating the onset of photosynthetic inactivation and freeze dehydration. (*c*) Representative *T-F*_0_ curve for *CT*_MAX_ (inflection point is the *T*_crit_) where leaf sample temperature (°C) increases beyond tolerance thresholds where the leaf rapidly emits more fluorescence (*F*_0_, relative units), indicating the onset of photosynthetic inactivation and potential damage. The example *T-F*_0_ curve for (*b*) *CT*_MIN_ is derived from a leaf sample on dry filter paper cooled at 15°C h^-1^ and for (*c*) *CT*_MAX_ is derived from a leaf sample on dry filter paper heated at 30°C h^-1^. The direction of arrows below the *x*-axes indicates the direction of temperature change.

We used a Pulse Amplitude Modulated (PAM) chlorophyll fluorescence imaging system (Maxi-Imaging-PAM; Heinz Walz GmbH, Effeltrich, Germany) mounted 185 mm above the Peltier plate (imaging area of approximately 120 × 90 mm) to measure fluorescence parameters. A weak blue pulse modulated measuring light (0.5 μmol photons m^-2^ s^-1^) was applied continuously at low frequency (1 Hz) to measure basal chlorophyll fluorescence (*F*_0_) from the LHCII without driving PSII photochemistry. A red Perspex hood filtered ambient light from the samples and the camera, and the entire Maxi-Imaging-PAM assembly was covered by thick black fabric so that all measurements were made in darkness. Leaves were dark adapted for 30 minutes to oxidise all PSII acceptors and obtain the basal *F*_0_ values and then a single saturating pulse at 10,000 μmol photons m^-2^ s^-1^ was applied for 720 ms to determine the maximal fluorescence (*F*_M_) when the photosystem reaction centres are closed. Variable fluorescence (*F*_V_) was calculated as *F*_M_ – *F*_0_ and the relative maximum quantum yield of PSII photochemistry (*F*_V_/*F*_M_) was derived. *F*_V_/*F*_M_ is frequently used as a rapid measurement of stress or relative health of leaves, where optimal *F*_V_/*F*_M_ values of non-stressed leaves are around 0.83 (Baker 2008; Murchie and Lawson 2013). Because our intention was to compare methods, we aimed for a uniform sample of leaves, and therefore we used *F*_V_/*F*_M_ values > 0.65 to subset data to exclude any damaged leaves and focus on the *T-F*_0_ of only healthy leaves. This conservative sample exclusion process resulted in some experimental conditions or species with uneven and lower sample sizes.

In each assay, we selected circular areas of interest that were as large as could fit within the boundaries of each leaf using the Maxi-Imaging-PAM software, such that the *F*_0_ values were measured on the widest part of each leaf. One minute after measuring *F*_V_/*F*_M_, the heating/cooling program was started simultaneously with the continuous recording of *F*_0_ values at set intervals with specifics varying depending on duration of the assay reflecting memory capacity limits of the Maxi-Imaging-PAM (see below). For hot *T-F*_0_ measurements, the initial set temperature held for dark adaptation of the leaves and *F*_V_/*F*_M_ was 20°C, which was then heated to 60°C at varying rates (see heating/cooling rate experiment). For cold *T-F*_0_ measurements, the assays were conducted in a cold room (set temperature: 4 ± 2°C) so that the Peltier plate could reach –20°C. At ambient room temperatures of ∼20–22°C, the Peltier plate can reach approximately –14°C before the plate heat output restrains cooling capacity. The initial set temperature held for dark adaptation of the leaves and *F*_V_/*F*_M_ was 4°C, which was then cooled down to –20°C.

The *T-F*_0_ curve produced by heating/cooling the Peltier plate (and leaf samples) is characterised by a stable or slow-rise in *F*_0_ values until a critical temperature threshold where there is a fast rise in *F*_0_. With temperature on the *x*-axis and *F*_0_ on the *y*-axis, the inflection point of extrapolated regression lines for each of the slow and fast rise phases of the temperature-dependent chlorophyll fluorescence response is the critical temperature, *T*_crit_ (Knight and Ackerly 2002; Neuner and Pramsohler 2006). The term *T*_crit_ is ambiguous outside of this context when both hot and cold thermal tolerance assays are conducted within the same study. Hereafter, we refer to *T*_crit_ only as the temperature extrapolated at the inflection point, and elsewhere use accepted nomenclature used in thermal biology, *CT*_MAX_ and *CT*_MIN_, as upper (heat) and lower (cold) thermal limits of leaf thermal tolerance (e.g., Sinclair *et al*. 2016; Janion-Scheepers *et al*. 2018). Figure 1 presents representative *T-F*_0_ curves and the calculations of *T*_crit_ values for freezing leaves, where the fast rise phase occurs abruptly (Fig. 1*b*), and for heating leaves where the fast rise phase is relatively gradual (Fig. 1*c*). The inflection point was calculated using a break-point regression analysis of the mean leaf temperature estimated from two thermocouples attached to leaves on the plate and relative *F*_0_ values using the *segmented* R package (Muggeo 2017) using the R Environment for Statistical Computing (R Core Team 2020). We provide example files and example R code for extracting *T*_crit_ values from *T-F*_0_ curves at https://github.com/pieterarnold/Tcrit-extraction.

### Surface wetness experiment: effect of wet vs dry surfaces for leaves on CT_MIN_ and CT_MAX_

Most experiments that measure *T-F*_0_ have measured leaf samples with all excess surface moisture removed, on a dry surface. However, maintaining water content of detached leaves by providing a wet surface where leaves were placed on top could be a viable way to facilitate water uptake and keep leaf samples hydrated. In our experiment, leaves were placed on a filter paper surface. For the wet surface treatment, leaves were placed as described above and then the filter paper was saturated with MilliQ water-soaked paper towels with excess water absorbed with dry paper towel thereafter. We compared *T-F*_0_ curves and *T*_crit_ estimates for both heat and cold tolerance assays at a heating/cooling rate of 60°C h^-1^ where leaves were placed on top of either wet or dry filter paper surfaces. A small subset of leaves on wet and dry surfaces were also measured for *CT*_MIN_ and *NT* at 15°C h^-1^ in addition to the 60°C h^-1^ experiment.

### Heating/cooling rate experiment: effect of heating/cooling rate on CT_MAX_ and CT_MIN_

Studies on thermal tolerance limits vary substantially in their set heating/cooling rate (Table S1), ranging from 30 to > 600°C h^-1^ in studies on heat tolerance limits (*CT*_MAX_) and from 1 to 10°C h^-1^ in studies on cold or freezing tolerance limits (*CT*_MIN_). The difference in magnitude between heat and cold tolerance limits reflects differences in natural potential rates of heating and cooling, where leaves may rapidly increase in temperature (> 240°C h^-1^ for a short period (Vogel 2009)) but cooling occurs far more slowly (rarely exceeding 5°C h^-1^ (Buchner and Neuner 2009)). It stands to reason that the more than 10-fold difference in heating or cooling rates used among studies would affect the estimates and thus comparability of *T*_crit_, but this effect is not well understood. We chose a wide range of heating/cooling rates for both hot and cold with the aim to determine how the *T*_crit_ estimate for *CT*_MIN_ and *CT*_MAX_ changes with heating/cooling rate. We compared *T-F*_0_ curves and *T*_crit_ estimates from different heating/cooling rates for both heat (6, 15, 30, 45, 60, 120, 240°C h^-1^) and cold (3, 6, 15, 30, 60, 240°C h^-1^) tolerance assays where the filter paper was dry, and measurements were conducted in darkness. For 240, 60, and 30°C h^-1^ heating/cooling rates, *F*_0_ was recorded at 10 s intervals, 20 s for 15 and 6°C h^-1^ heating/cooling rates, and 30 s for 3°C h^-1^ heating/cooling rates due to the 1000 record limit after which the Maxi-Imaging-PAM software stops recording.

### Statistical analyses

The dataset was trimmed by removing leaves that had initial *F*_V_/*F*_M_ values below 0.65, which was a value chosen to identify and remove unhealthy or damaged leaves, hence sample sizes varied among species and experimental conditions. Summary data (mean ± standard error) is reported in Table S2. Data that matched conditions used in all experiments were used for multiple analyses (e.g., hot assay, heating/cooling rate of 60°C h^-1^, dry filter paper could be used for all). Linear regression models were implemented using the *stats* package in the R environment for statistical and graphical computing (v3.5.1) (R Core Team 2020). Models were specified with *CT*_MIN_ or *CT*_MAX_ as the response variable and fixed categorical predictors of either wet/dry or heating/cooling rate depending on the experiment. *F*_V_/*F*_M_ was always included as a fixed covariate. We first fit models combining the three species for a given experiment, and then we fit species-specific models. Preliminary models were linear mixed effects regression models that included individual plant as a random factor, but in almost all cases, the term explained essentially zero variance, so we removed the random term in favour of a simpler linear model. Tables report model parameter estimates with statistical significance at *p* < 0.05 indicated in bold and with * symbols. Supplementary tables (Tables S3–S6) report full statistical model output. Figures show means with non-parametric bootstrapped 95% confidence intervals (95% CIs) derived from the *Hmisc* R package (Harrell 2019). Finally, predicted temperature threshold estimates were modelled as a quadratic function of heating/cooling rate treated as a continuous variable for visualisation purposes. The data that support the findings of this study are openly available in the figshare repository: 10.6084/m9.figshare.12545093.

## Results

### Overview

The Peltier plate and chlorophyll fluorescence Maxi-Imaging-PAM system allows us to measure *T-F*_0_ (Fig. 1) on many leaves simultaneously. In these experiments, we measured up to 30 whole leaf samples in a single experimental run, which could take as little as 90 minutes including dark adaptation, leaf set up on the surface, and the temperature heating/cooling rate (at 60°C h^-1^). The Peltier plate can easily accommodate a much greater number of smaller leaves, leaf discs, or leaf sections for even higher throughput phenotyping if required (Fig. S1).

### Surface wetness experiment: effect of wet vs dry surface for leaves on CT_MIN_ and CT_MAX_

The effect of water saturating the filter paper was clearly apparent for *T*_crit_ value estimates for *CT*_MIN_ (Fig. 2*a*) but not *CT*_MAX_ (Fig. 2*b*). For all species combined and when the three species were analysed separately, *CT*_MIN_ values were significantly and consistently less negative (less cold tolerant) for leaves on wet surfaces than on dry ones, by 3–4°C (Table 1, S3, Fig. 2*a*). Variation in *CT*_MIN_ was independent of the initial *F*_V_/*F*_M_ of leaves. The *CT*_MAX_ of leaves with a wet paper surface did not differ significantly from dry ones both among and within species (all *p* > 0.2; Table 1, S3, Fig. 2*b*), although the three species had different *CT*_MAX_ estimates. Leaves with higher *F*_V_/*F*_M_ had higher *CT*_MAX_ for *W. ceracea*.

**Table 1.** Summary of analyses of all species and species-specific effects of wet *vs* dry filter paper surface on *CT*_MIN_ and *CT*_MAX_.

| **Response: $CT_{\text{MIN}}$** | **All species** | ***W. ceracea*** | ***M. citrina*** | ***E. rubra*** |
| --- | --- | --- | --- | --- |
| **Fixed effects** | **Estimate** | **Estimate** | **Estimate** | **Estimate** |
| Dry surface / *E. rubra* (intercept) | **-18.36\*** | Intercept: -5.71 | Intercept: **-20.36\*** | Intercept: **-31.26\*\*** |
| Wet surface | **3.81\*\*\*** | **3.92\*\*\*** | **2.98\*\*** | **3.99\*\*\*** |
| $F_V/F_M$ | 6.19 | -9.89 | 4.72 | 23.54 |
| *M. citrina* | **-3.50\*\*\*** | -- | -- | -- |
| *W. ceracea* | -0.42 | -- | -- | -- |
| $R^2$ | 0.464 | 0.288 | 0.374 | 0.527 |

| **Response: $CT_{\text{MAX}}$** | **All species** | ***W. ceracea*** | ***M. citrina*** | ***Q. phellos*** |
| --- | --- | --- | --- | --- |
| **Fixed effects** | **Estimate** | **Estimate** | **Estimate** | **Estimate** |
| Dry surface / *M. citrina* (intercept) | **32.76\*\*\*** | Intercept: 6.90 | Intercept: **36.31\*** | Intercept: **47.34\*\*\*** |
| Wet surface | -0.55 | -1.47 | 0.32 | -0.63 |
| $F_V/F_M$ | 18.20 | **46.02\*** | 13.16 | 2.32 |
| *Q. phellos* | **2.01\*\*** | -- | -- | -- |
| *W. ceracea* | **-4.47\*\*\*** | -- | -- | -- |
| $R^2$ | 0.593 | 0.213 | 0.028 | 0.041 |

**Fig 2.**
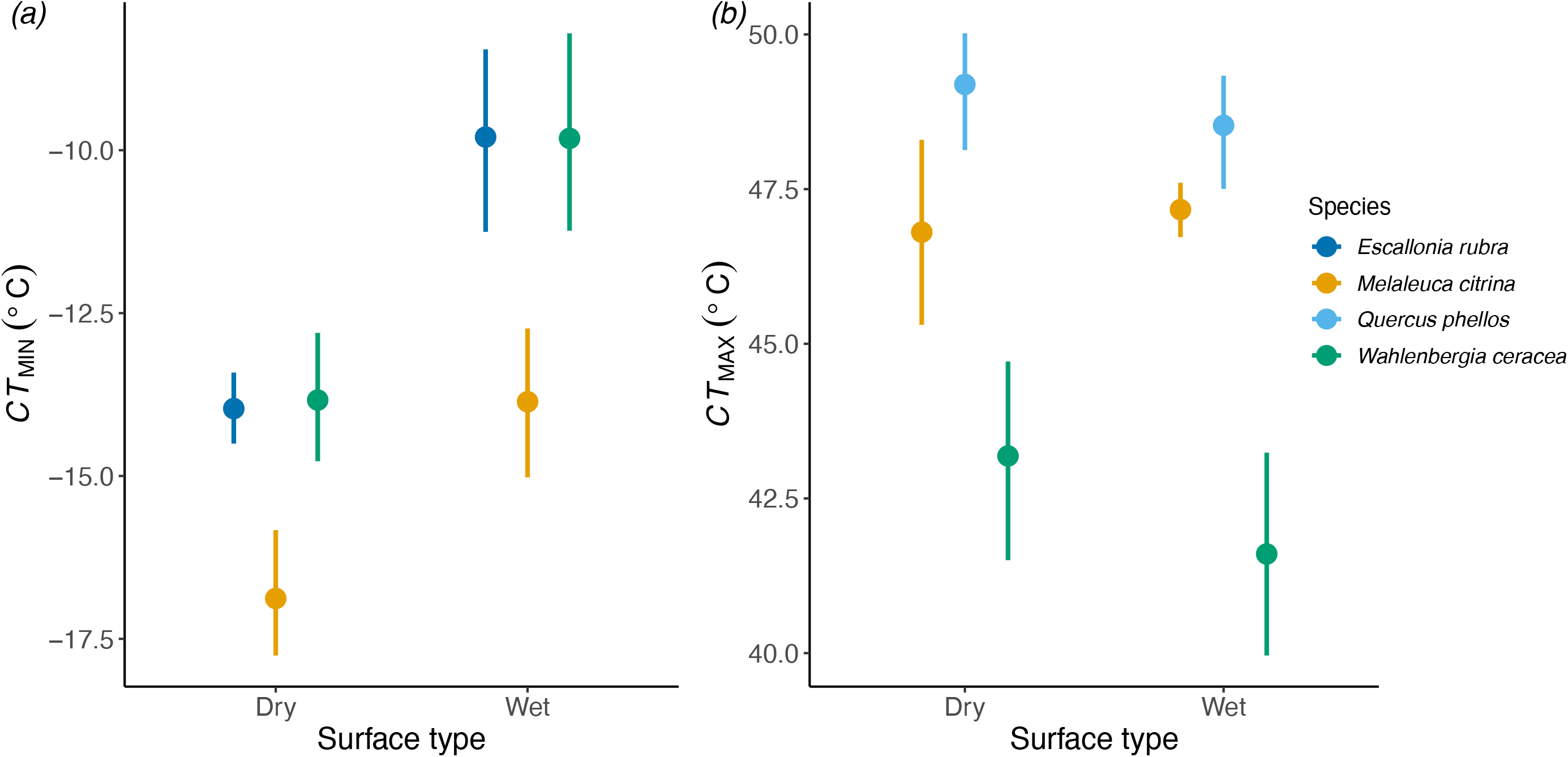
The effect of varying surfaces (dry *vs* wet filter paper) on the *CT*_MIN_ and *CT*_MAX_ estimates (°C) from basal chlorophyll fluorescence (*F*_0_, relative units) of leaves. We tested how (*a*) *CT*_MIN_ and (*b*) *CT*_MAX_ estimates of leaves from four plant species under standard dry conditions (dry filter paper surface) differed from wet conditions (wet filter paper surface). All estimated were obtained using a standard heating/cooling rate of 60°C h^-1^. Data points are means and 95% CIs that overlay raw data (*n* = 12–25 per treatment × species combination).

### Surface wetness × heating/cooling rate experiment: effects on CT_MIN_ and NT

*CT*_MIN_ of leaves of all species was higher on a wet surface and generally lower at faster cooling rates compared to leaves on a dry surface at slower cooling rate (Table S4). However, the interaction between surface wetness and cooling rate never had a significant effect on *CT*_MIN_; leaves on a wet surface had a consistently higher *CT*_MIN_ than those on a dry surface at both 15 and 60°C h^-1^. A small subset of 17 leaves could be used to test whether surface wetness and cooling rates affected *NT*, however, due to this low sample size, we opted not to formally analyse these data, but present descriptive findings in Fig. S2. *NT* of leaves measured on a wet surface occurred at higher temperatures (around –7°C) independently of cooling rate, however *NT* occurred at lower temperatures on leaves on a dry surface, and perhaps slightly lower on leaves exposed to a faster cooling rate (Fig. S2). *NT* generally occurred at temperatures 2–4°C higher than *CT*_MIN_, and the mean difference between *CT*_MIN_ and *NT* was 1°C lower on a wet surface compared to a dry surface (Fig. S2).

### Heating/cooling rate experiment: effect of heating/cooling rate on CT_MAX_ and CT_MIN_

Varying heating/cooling rate affected the estimate of *T*_crit_ for *CT*_MIN_ and *CT*_MAX_ considerably, however each species responded differently. For *CT*_MIN_, slow cooling rates (< 10°C h^-1^) are standard practice and here we used 3°C h^-1^ as the reference category. We found no significant differences between 3, 6, 15, or 30°C h^-1^ cooling rates overall, but when the plate was cooled at faster rates, the *CT*_MIN_ values became very different to the slower cooling rates. At 60 and 240°C h^-1^ *CT*_MIN_ was significantly lower relative to 3°C h^-1^ for *M. citrina* and *E. rubra* (Table 2, S5). For *M. citrina*, the values shifted depending on cooling rate, but with no clear pattern (Fig. 3*a*). In contrast, *E. rubra* had stable *CT*_MIN_ values for 3, 6, and 15°C h^-1^ and more negative values as cooling rate increased to 30, 60, and 240°C h^-1^ (Table 2, S5, Fig. 3*a*). *CT*_MIN_ for *W. ceracea* was similar across most cooling rates and was only significantly different from when the cooling rate was 30°C h^-1^ (Table 2, S5). Variation in *CT*_MIN_ was independent of the initial *F*_V_/*F*_M_ of leaves.

**Table 2:** Summary of analyses of all species and species-specific effects of variable temperature heating/cooling rate on *CT*_MIN_ and *CT*_MAX_.

| **Response: $CT_{\text{MIN}}$** | **All species** | ***W. ceracea*** | ***M. citrina*** | ***E. rubra*** |
| --- | --- | --- | --- | --- |
| **Fixed effects** | **Estimate** | **Estimate** | **Estimate** | **Estimate** |
| Cooling rate = 3°C h<sup>-1</sup> / *E. rubra* (Intercept) | **-11.38\*\*\*** | Intercept: **-40.89\*\*** | Intercept: **-16.82\*\*\*** | Intercept: **-11.58\*\*** |
| Cooling rate = 6°C h<sup>-1</sup> | -0.33 | 0.62 | **-1.81\*** | -0.15 |
| Cooling rate = 15°C h<sup>-1</sup> | -0.32 | -0.12 | -0.80 | -0.10 |
| Cooling rate = 30°C h<sup>-1</sup> | 0.75 | **1.67\*\*** | 0.91 | -0.74 |
| Cooling rate = 60°C h<sup>-1</sup> | **-1.34\*\*** | 0.75 | **-3.51\*\*\*** | **-2.47\*\*\*** |
| Cooling rate = 240°C h<sup>-1</sup> | -0.80 | 0.70 | **-1.74\*** | **-1.90\*\*** |
| $F_V/F_M$ | -0.89 | 32.04 | 4.66 | 0.12 |
| *M. citrina* | **-2.18\*\*\*** | -- | -- | -- |
| *W. ceracea* | **-1.53\*\*** | -- | -- | -- |
| Marginal R<sup>2</sup> | 0.230 | 0.126 | 0.332 | 0.220 |

| **Response: $CT_{\text{MAX}}$** | **All species** | ***W. ceracea*** | ***M. citrina*** | ***E. rubra*** |
| --- | --- | --- | --- | --- |
| **Fixed effects** | **Estimate** | **Estimate** | **Estimate** | **Estimate** |
| Heating rate = 60°C h<sup>-1</sup> / *E. rubra* (Intercept) | **27.79\*\*\*** | Intercept: 14.87 | Intercept: **27.79\*\*\*** | Intercept: **41.75\*** |
| Heating rate = 6°C h<sup>-1</sup> | -0.68 | **1.60\*** | **-7.71\*\*\*** | 1.38 |
| Heating rate = 15°C h<sup>-1</sup> | **2.43\*\*\*** | **-2.00\*\*** | **-4.68\*\*\*** | -1.40 |
| Heating rate = 30°C h<sup>-1</sup> | **-1.74\*\*** | **-2.11\*\*\*** | **-2.10\*\*** | **-1.31\*** |
| Heating rate = 45°C h<sup>-1</sup> | **-1.48\*\*** | -0.72 | -0.95 | **-2.68\*\*\*** |
| Heating rate = 120°C h<sup>-1</sup> | 1.00 | **1.78\*\*** | **2.24\*** | -0.45 |
| Heating rate = 240°C h<sup>-1</sup> | **2.03\*\*\*** | **2.78\*\*\*** | **3.76\*\*\*** | -0.13 |
| $F_V/F_M$ | **23.79\*\*\*** | **38.48\*\*\*** | **21.98\*\*** | 5.15 |
| *M. citrina* | **-1.30\*\*\*** | -- | -- | -- |
| *W. ceracea* | **-1.24\*\*** | -- | -- | -- |
| Marginal R<sup>2</sup> | 0.429 | 0.619 | 0.863 | 0.319 |

Bold indicates significance at $p < 0.05$ , \*: $p < 0.05$ , \*\*: $p < 0.01$ , \*\*\*: $p < 0.001$ . Intercepts marked as significant are different from zero. Full statistical reporting is provided in Tables S5 and S6.

**Fig 3.**
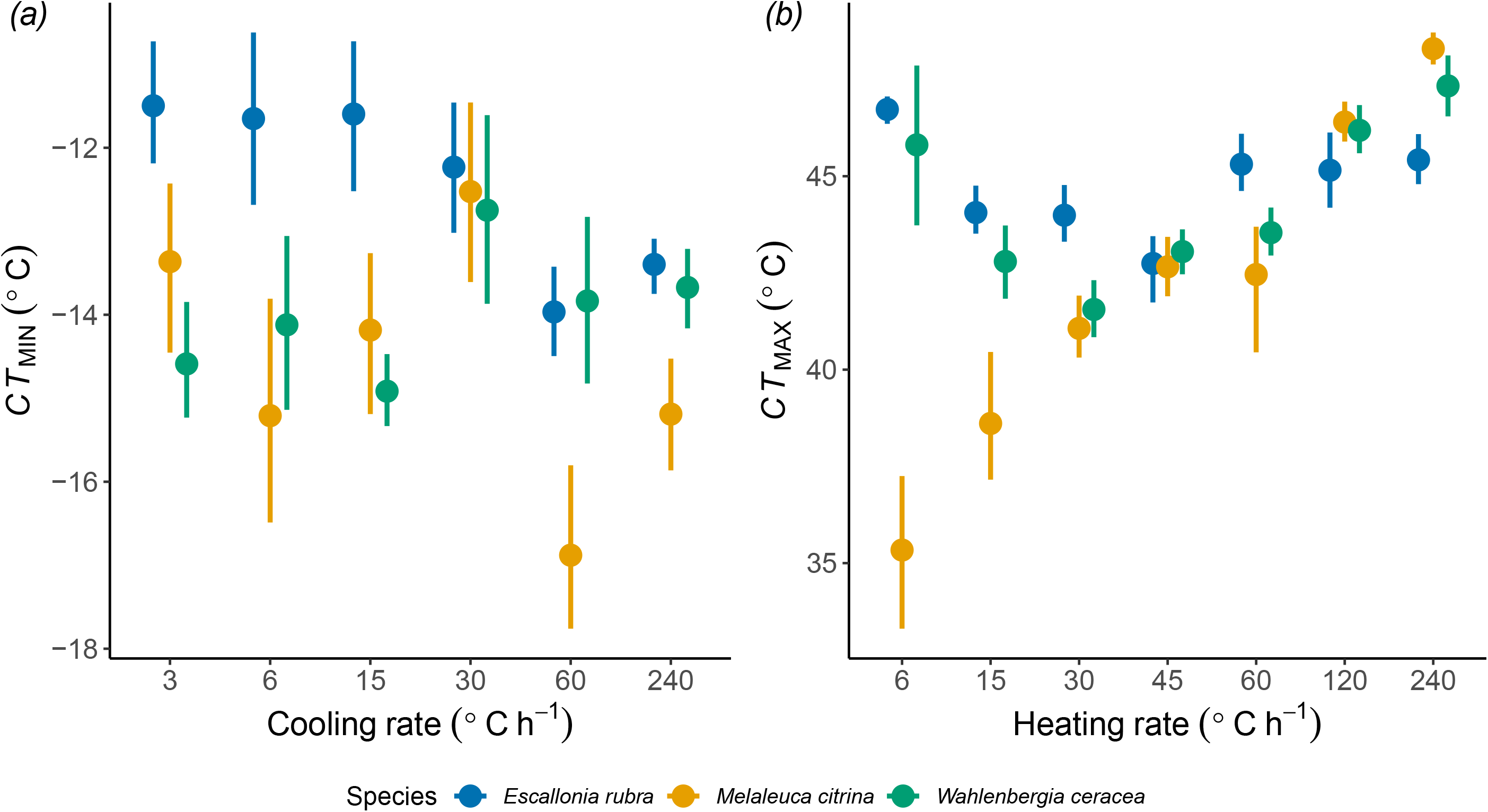
The effect of varying heating/cooling rate (°C h^-1^) on the *CT*_MIN_ and *CT*_MAX_ estimates (°C) from basal chlorophyll fluorescence (*F*_0_, relative units) of leaves. We tested how (*a*) *CT*_MIN_ and (*b*) *CT*_MAX_ estimates of leaves from three plant species were affected by changing the temperature stress at different heating/cooling rates. Data points are means and 95% CIs that overlay raw data (*n* = 6–20 per treatment × species combination).

*CT*_MAX_ is typically measured with a heating rate of 60°C h^-1^, so this was used as a reference against which all other heating rates were compared. *CT*_MAX_ was highly dependent on heating rate, where rates slower than 60°C h^-1^ produced significantly lower *CT*_MAX_ estimates, except for 6°C h^-1^. Heating rates higher than 60°C h^-1^ resulted in higher *CT*_MAX_ estimates, significantly so for 240°C h^-1^ but not 120°C h^-1^ (Table 2, S6). However, stark species-specific responses were evident. *CT*_MAX_ in *M. citrina* was very low at heating rates of 6 and 15°C h^-1^ and increased significantly and consistently with faster heating rates: only 45 and 60°C h^-1^ yielded similar *CT*_MAX_ values (Table 2, S6, Fig. 3*b*). In contrast, *CT*_MAX_ in *E. rubra* was higher at the slowest rate (although the effect was marginal) compared to 60°C h^-1^ but significantly lower at 30 and 45°C h^-1^ and not different from 120 and 240°C h^-1^ (Table 2, S6, Fig. 3*b*). Similarly, *W. ceracea* had significantly higher *CT*_MAX_ values at 6°C h^-1^, but also at 120 and 240°C h^-1^. Only 45 and 60°C h^-1^ produced *CT*_MAX_ values for *W. ceracea* that were not significantly different (Table 2, S6, Fig. 3*b*). In all analyses except *E. rubra* individually, *F*_V_/*F*_M_ had a significant positive relationship with *CT*_MAX_.

### Heating/cooling rate experiment: predicted thermal limits as a function of heating/cooling rate

We then modelled predicted *CT*_MAX_ and *CT*_MIN_ values against heating/cooling rate as a continuous variable using a quadratic function to visualise the interspecific differences in response to different heating/cooling rates when measuring thermal limits (Fig. 4*a, b*). The difference between 60 and 240°C h^-1^ introduced extreme uncertainty in the predicted *CT*_MIN_ for *M. citrina*, so the 240°C h^-1^ rate was removed from the visualisation. The shape of each species’ *CT*_MAX_ and *CT*_MIN_ response to heating/cooling rate were clearly distinct from one another and only *E. rubra* had a relatively stable predicted *CT*_MAX_ value across all measured heating/cooling rates. The variance tends to increase with faster heating/cooling rates for *CT*_MIN_, but the pattern is less clear for *CT*_MAX_.

**Fig 4.**
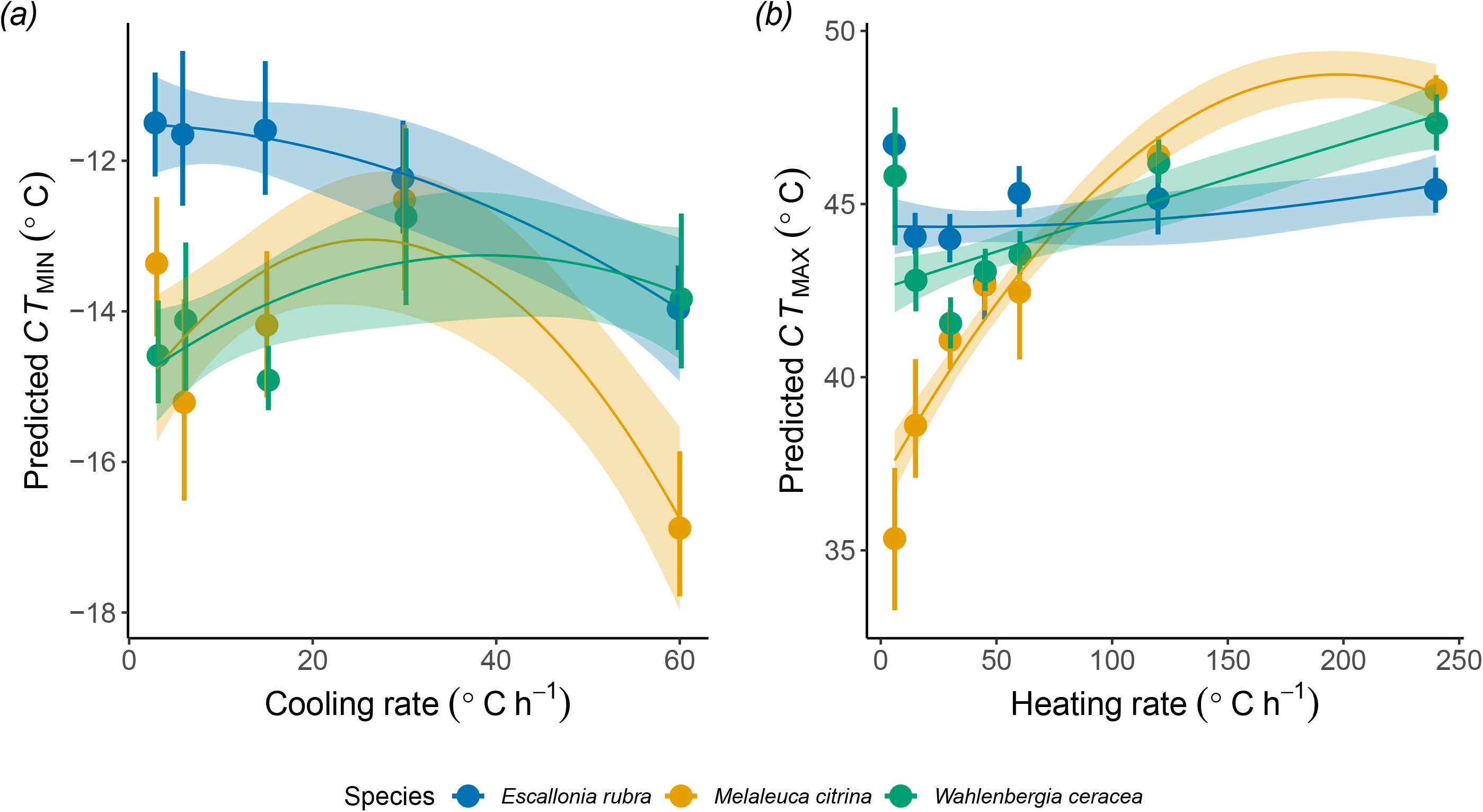
The effect of heating/cooling rate (°C h^-1^) as a continuous variable on the (*a*) predicted *CT*_MIN_ and (*b*) predicted *CT*_MAX_ estimates (°C) in leaves from three plant species. Data points are means and 95% CIs (*n* = 6–20 per treatment × species combination) with predicted response curves modelled with quadratic functions separately for each species.

## Discussion

We sought to develop a reliable, high-throughput method for assessing thermal tolerance limits of the photosynthetic apparatus. Many methods are used for measuring plant thermal tolerance limits, but such variation has potential consequences for generating reasonable interpretations and interspecific comparisons. Often, the rationale behind a published method is unclear and the impacts of small methodological differences are difficult to assess (Geange *et al*. 2021). To address this, we have demonstrated a method for measuring both cold and heat tolerance limits of leaves using a thermoelectric plate and chlorophyll imaging fluorescence. In line with previous applications of this technique, we provide evidence for the effects of controllable experimental variables on estimates of *CT*_MIN_ and *CT*_MAX_. We quantify the significant effects of measurement conditions and show that using a wet *vs* dry surface for measuring *CT*_MIN_ and that variation in heating/cooling rates leads to substantial differences in *CT*_MIN_ and *CT*_MAX_. We aimed to develop a practical method that maximises informative value and minimises experimental noise among samples. In the case of heating/cooling rate, there is high species specificity. Below we outline potential mechanistic explanations for our findings along with testable hypotheses, and then propose best practices for measuring the thermal tolerance limits of leaves.

### Pros and cons of the T-F_0_ Peltier plate-Maxi-Imaging fluorimeter method

Measuring the temperature-dependent change in basal chlorophyll fluorescence is one of several potential methods that researchers can use to quantify the critical thermal limits of photosynthesis activation and photosynthetic apparatus stability (Ilík *et al*. 2003). The method that we present here offers improvements over earlier and alternative versions that use bulky water baths or freezing chambers, or smaller capacity Peltier plates (e.g., Schreiber and Berry 1977; Braun *et al*. 2002; Knight and Ackerly 2002; Neuner and Pramsohler 2006), and adds several key features. The Peltier plate-Maxi-Imaging fluorimeter system is relatively compact and transportable for field applications when provided with a continuous power source. It offers precise temperature control (± 0.1°C precision and ± 1°C tolerance) and high versatility by being programmable for both cooling and heating rapidly at set rates. It can be programmed for stepwise temperature treatments or non-linear temperature programs, or temperature shock treatments depending on the desired application. Furthermore, the *T-F*_0_ curve allows for the calculation of other parameters (e.g., Knight and Ackerly 2002), including the temperatures at 50% or 100% of relative *F*_0_ (*T*_50_ and *T*_max_, respectively) and ice nucleation temperatures (*NT*) for cold tolerance assays if each leaf sample has a thermocouple attached to it (e.g., Briceño *et al*. 2014). When using detached leaves or leaf discs, the potential throughput of the system is substantial (Fig. S1). The 120 × 90 mm optimal imaging area on the Peltier plate can fit > 100 leaf discs or small leaf samples up to 1 cm^2^ or > 30 samples that are up to 2 cm^2^ each, thus throughput is mostly constrained by sampling and setting up that many leaves.

As with any laboratory equipment, there are limitations to the Peltier plate-Maxi-Imaging fluorimeter system. Unlike freezing chambers, this system does not allow for whole-plant measurements. There is some software modification required for controlling the heating/cooling rates using the Peltier plate system, although newer temperature controllers and software revisions than those used here are now available. The Peltier plate-Maxi-Imaging fluorimeter system is a versatile phenotyping tool for thermal tolerance, ecophysiology, and photosynthesis research. Below, we discuss the results of testing the system with wet and dry filter paper as surfaces and the effects of heating/cooling rates.

### A dry surface avoids experimental artefacts

Using wet filter paper as a surface for the leaf samples significantly reduced the apparent measured *CT*_MIN_ but had no effect on *CT*_MAX_. Wet filter paper was initially tested to attempt to avoid leaf dehydration by providing a saturating atmosphere, preventing leaf evapotranspiration. In our cold tolerance assay, freezing of the water in the wet filter paper most likely began propagating ice from outside the leaf into the apoplastic space, thereby freezing the apoplast in the leaf tissue at higher temperatures than leaves on the dry surface. When radiative frost occurs, air humidity condenses on the leaf surface, resulting in a wet leaf surface that may induce heterogenous extrinsic nucleation in natural frosts (Pearce 2001). Thus, the wet filter paper surface acted as an extrinsic ice nucleator and likely prevented the leaves from supercooling (Sakai and Larcher 1987; Pearce 2001; Larcher 2003). Our exploratory tests between wet and dry surfaces at different cooling rates demonstrated that on a dry filter paper surface, leaves appeared to supercool 2–4°C below those leaves on a wet surface. *NT* occurred earlier and at temperatures closer to *CT*_MIN_ on the wet surface and was more variable in comparison to leaves on a dry surface. Although this supercooling phenomenon requires further targeted investigation in future, our initial tests suggest that a wet surface induces earlier ice formation and propagation at warmer temperatures and hence reduces leaf supercooling capacity, and that supercooling capacity might be exacerbated by faster cooling rates.

The initial water status of leaf samples is still crucial, as water-stressed leaves can have compromised (Verslues *et al*. 2006) or even enhanced stress tolerance (Havaux 1992). Therefore, we recommend that detached leaves should be transported in a manner that maintains leaf water content after sampling (e.g., sealing leaves with plastic film wrap, using damp paper towel, or cut stems placed in water) so that leaves are either maintained at collection conditions or fully hydrated at the start of the thermal tolerance assay.

### Maximising throughput without compromising results

A wide range of heating/cooling rates have been used in previous studies of thermal limits to photosynthesis (Table S1). We have demonstrated that heating/cooling rate strongly influences both *CT*_MIN_ and *CT*_MAX_ values with varying magnitude and complex patterns for different species. Indeed, we saw such strong species-specific responses to different heating/cooling rates (particularly for heat) that if one were to measure the *CT*_MAX_ for three species measured at the same heating rate of 45°C h^-1^, they would conclude that all the species have identical heat threshold temperatures, yet the same experiment conducted with a heating rate of 6°C h^-1^ and 240°C h^-1^ would result in entirely different, and opposing, conclusions. For comparative studies that measure species with different leaf morphology, physiology, and biochemical constituents, it is crucial that we clarify and refine what physiological event(s) we aim to characterise with the *T-F*_0_ approach. From a practical standpoint, our aim was to identify the fastest heating/cooling rates that would allow repeatable, interpretable measures of *T*_crit_.

Heating rates will determine the potential for activation and extent of the upregulation of physiological processes and protective mechanisms within the leaf when approaching thermal extremes (Bilger *et al*. 1984; Frolec *et al*. 2008). The rise in *F*_0_ during a measure of *CT*_MAX_ indicates when photosynthetic activity is markedly reduced and thereafter the thylakoid membrane is disrupted (Havaux *et al*. 1988; Nauš *et al*. 1992). If leaf samples are heated only up to the temperature of the initial rise in *F*_0_, *CT*_MAX_, and then cooled, it is possible that membrane disruption can be reversed (Yamane *et al*. 1997; Frolec *et al*. 2008). However, irreversible damage to PSII through physiological changes to the photosynthetic apparatus and then physical membrane separation (i.e., denaturation) is correlated with the continued rapid rise and maxima of *F*_0_ with sustained extreme temperatures (Terzaghi *et al*. 1989; Frolec *et al*. 2008). Specifically, the first peak in *F*_0_ shortly after *CT*_MAX_ and between 40–50°C is due to irreversible inactivation of PSII and the secondary *F*_0_ peak between 55–60°C originates from the denaturing of chlorophyll-containing protein complexes (Ilík *et al*. 2003). Leaves can reduce the photochemical and oxidative impairment induced by heat stress by thermal dissipation of excessive excitation energy to maintain PSII in an oxidative state, and by upregulating heat shock proteins and antioxidant activity (Allakhverdiev *et al*. 2008; Silva *et al*. 2010). Changes to the lipid composition of the thylakoid membrane reduces the fluidity of the membrane thereby being more stable at high temperatures (Allakhverdiev *et al*. 2008). The upregulation of these protective mechanisms of PSII can occur relatively quickly, sometimes < 1 h of heat stress (Havaux 1993), thus how protected the leaf is against PSII inactivation will depend on the heating rate.

For cold tolerance assays, cooling rates likely modify the dynamic and primary site of ice nucleation. Intrinsic ice nucleation may lead to ice formation in the xylem (Hacker and Neuner 2007), while extrinsic nucleation occurs at the leaf epidermis (Pearce and Ashworth 1992). Rates of cooling may also influence supercooling capacity; usually faster cooling (within the range of this study) increases supercooling capacity (Gokhale 1965). Despite most freezing studies using cooling rates that are more reminiscent of natural freezing rates (≤ 5°C h^-1^), we did not find a clear difference among *CT*_MIN_ values at cooling rates of 3, 6, and 15°C h^-1^. We hypothesise that reducing the temperature relatively slowly (e.g., ≤ 15°C h^-1^) could allow the cell to adjust osmotically and partially counterbalance the reduced water potential of the frozen apoplast restricting cell dehydration, which would be avoided at faster cooling speeds. Thus, the consideration for the freezing tolerance cooling rates becomes a question of what is the greatest cooling rate that allows more realistic osmotic adjustments within the leaf.

For *W. ceracea* and *E. rubra*, increasing temperature slowly (< 30°C h^-1^) appears to allow time for induction of protective mechanisms such that slower heating rates result in higher *CT*_MAX_ values. Conversely, changing temperature more quickly (30–60°C h^-1^) prevents membranes from inducing heat-hardening or for antioxidants to be upregulated and take effect, such that measured heat tolerance limits is relatively stable at these heating rates. Our results indicate that beyond a rate of 60°C h^-1^, the increase in *F*_0_ occurs more slowly than the temperature increase and the temperature of the leaf samples (as measured by thermocouples) also lags significantly behind the temperature of the Peltier plate, thus the *CT*_MAX_ may be overestimated (Fig. 3*b*). Hence, using the thermistor (plate) temperature will overestimate the temperature of the leaf, and therefore, its tolerance limit. Furthermore, the faster that the plate temperature is changed, the more potential variation among leaf temperatures. We acknowledge that the method could be improved by using individual thermocouples for each leaf sample, particularly for cold tolerance to measure ice nucleation temperature (*NT*), however, we have verified that there is minimal variation (±1°C) across the Peltier plate surface.

The species specificity of the heating rate dependence of *CT*_MAX_ was striking, particularly in the case of *M. citrina*. A slow heating rate of 6°C h^-1^ results in a very low estimate for *CT*_MAX_ of only 36°C, which suggests that the heat tolerance of this species is poor, yet at heating rates ≥ 30°C h^-1^, this species is apparently as or more heat tolerant than the other species. Slow heating rates mean that the leaves are slow to reach more stressful temperatures, but also that they are held at these temperatures for longer periods of time. We hypothesise that the lower heat tolerance limit at slow heating rates could be due to leaf water being tightly bound and preventing cooling via transpiration or the heated leaf oils being unable to volatilise, thereby destabilising membranes and effectively ‘slow-cooking’ the leaf. For this species, the higher heating rates are therefore likely more indicative of photosynthetic thermal tolerance limits.

The *T-F*_0_ method is a rapid measurement compared to other *F*_V_/*F*_M_-based assessments of thermal tolerance. Determining the temperature at which 50% of the potential thermal damage (lethal temperature) to the plant tissue occurs (*LT*_50_) is a common but very time-consuming technique that also requires more plant material. Different individual leaves are heated/cooled to and held at set temperatures for 1-3 h, and then *F*_V_/*F*_M_ is measured over 1-24 h post-thermal exposure to determine the point of irreversible damage. We note that *F*_0_ can be affected by leaf properties including the efficiency of PSII, the leaf chlorophyll content and ratios, and leaf thickness, which may affect thermal tolerance estimates more than those measured using *F*_V_/*F*_M_. Therefore, to better understand what occurs within a leaf during exposure to thermal extremes, it would be valuable to characterise the *T-F*_0_ curve and identify the *CT*_MIN_ and *CT*_MAX_ values for a plant. One could then heat/cool and hold leaf samples at these threshold temperatures for a set time, then measure *F*_V_/*F*_M_ with the same Maxi-Imaging fluorescence system to examine potential recovery from exposure to damaging temperatures (e.g., Buchner *et al*. 2015). Then, one could investigate the correlation between *CT* and *LT* metrics and determine the extent and reversibility of damage. A more complete micro-scale understanding of thermal tolerance responses and species specificity would be enhanced by exploring tissue biochemistry, the regulation of heat shock proteins, and gene expression at thermal extremes (Geange *et al*. 2021). At the macro end of the scale, remote sensing tools allows landscape scale estimations of photosynthetic tolerance to heating using the Photochemical Reflectance Index (PRI), which strongly relates to stress changes in photosynthetic machinery (Sukhova and Sukhov 2018; Yudina *et al*. 2020). Comparative studies on the accuracy and precision of different micro- and macro-scale techniques for estimating thermal tolerance of plants will be necessary for maximising agricultural and ecological monitoring efforts.

### Towards standardised approaches for comparative thermal tolerance research

There will never be a perfect one-size-fits-all method for comparative measures of plant photosynthetic thermal tolerance, but our exploration of method variation we find there is a reasonable set of conditions that will fit most. We advocate that researchers use well-hydrated leaves (unless hydration status is an element of their experiment) and dry surface for these measures. Doing so allows easy comparison across experiments and gives a more indicative measure of the lowest potential *CT*_MIN_.

We sought the maximum heating/cooling rate that was repeatable and reliable. Our results suggest that there is a point beyond which temperatures are changed too quickly and the *T*_crit_ value is exaggerated due to the change in *F*_0_ lagging the change in leaf temperature, especially in heat tolerance limit assays. For an experiment on a single or few species, pilot studies on the effects of heating/cooling rates are advisable. For broad interspecific studies, particularly in natural systems where other variables such as thermal history and the environment cannot be controlled, using a common rate for heating and for cooling is the only feasible approach. For such comparative work, we recommend a heating rate of not less than 30°C h^-1^ (up to 60°C h^-1^ to avoid any potential heat hardening) for *CT*_MAX_ and a cooling rate at or below 15°C h^-1^ for *CT*_MIN_. We recognise that this is a slower heating rate than often used for *CT*_MAX_ and a faster than usual cooling rate for *CT*_MIN_. However, we found that the 15°C h^-1^ rate was not significantly different to slower rates for *CT*_MIN_ and thus represents the most efficient rate that could yield results reflective of natural scenarios. For *CT*_MAX_, we argue that the 30– 60°C h^-1^ rates enable physiological mechanisms that would normally provide some thermal protection to the photosystem and cell membranes to be induced, without lag exaggerating *CT*_MAX_, and may therefore be a more realistic or relevant measurement of thermal tolerance than that provided by faster rates. These rates remain practical for achieving high throughput, especially with sample sizes that can be accommodated by large Peltier plates combined with the multi-sample imaging of Maxi-Imaging fluorimeters.

Clearly, any experimental thermal tolerance assay cannot perfectly mirror the conditions of a natural extreme thermal event. Rates of heating and cooling of plant tissues in nature are non-linear, not sustained, and strongly mediated by external conditions such as wind, solar radiation, season, and elevation (Sakai and Larcher 1987; Leuning and Cremer 1988; Vogel 2009). The researcher must always remain appreciative of how extrinsic factors could affect these values and interpretations thereof for their study system. However, *T-F*_0_ curves and derived *T*_crit_ values can indicate what the *potential* thermal limits of leaves are, under absolute conditions. The method provides power for comparative research, and also ample opportunity to explore the underlying mechanisms of species level differentiation. Moving toward a deeper understanding of the physiological processes conferring thermal tolerance is crucial in the changing climate where extreme weather events are increasing in frequency and intensity (Buckley and Huey 2016; Harris *et al*. 2018).

## Conclusions

The Peltier plate-Maxi-Imaging fluorimeter system described and tested here allows relatively high-throughput measurement of *T-F*_0_ and the critical thermal limits to inactivation of photosynthesis. This system offers great flexibility and substantially expands on previous versions. We have demonstrated that use of wet *vs* dry surface can significantly affect the *CT*_MIN_ estimate, but not *CT*_MAX_, and that heating/cooling rates have strong species-specific effects on both *CT*_MIN_ and *CT*_MAX_. Awareness of the physiological processes that underlie the rapid rise in *F*_0_ and consideration of interspecific differences in leaf physiology and biochemistry are essential for making effective choices in the rate of heating or cooling leaf samples. We recommend the use of parameters that maximise repeatability and efficiency of the measurements without introducing artefacts of heating/cooling rate. As plants around the world are exposed to more thermal extremes by the effects of climate change, versatile ecophysiological tools such as this Peltier plate-Maxi-Imaging fluorimeter system will be valuable for generating new insights in plant responses and thermal tolerance limits.

## Acknowledgements

We sincerely thank Ya Zhang for modifying the LabVIEW software for heating/cooling rate control, ANU plant services staff for maintaining glasshouse plants, and Jack Egerton and ANU workshop staff for technical support. We thank three anonymous reviewers and Loeske Kruuk for their constructive feedback on earlier versions of this manuscript. This research was supported by the Australian Research Council (DP170101681).

## Author contribution statement

PAA, KMG, AAC, and ABN designed the experiments. PAA, KMG, AAC performed the experiments and collected the data. PAA curated the data and performed the data analyses and visualisation. PAA, VFB, LAB, and ABN interpreted the results and wrote the manuscript with input from all authors.

## Supplemental material

The following supplemental materials are available.

**Supplemental Table S1:** Samples of heating/cooling rate variation from the literature.

**Supplemental Table S2:** Mean values for *CT*_MIN_, *CT*_MAX_, and *F*_V_/*F*_M_ for each species and experimental condition.

**Supplemental Table S3:** Full statistical reporting for effects of wet *vs* dry surface for *CT*_MIN_ and *CT*_MAX_.

**Supplemental Table S4:** Full statistical reporting for effects of wet *vs* dry surface combined with heating/cooling rate on *CT*_MIN_.

**Supplemental Table S5:** Full statistical reporting for effects of heating/cooling rate for *CT*_MIN_.

**Supplemental Table S6:** Full statistical reporting for effects of heating/cooling rate for *CT*_MAX_.

**Supplemental Figure S1:** Various experimental applications of the Peltier plate and chlorophyll fluorescence Maxi-Imaging-PAM system.

**Supplemental Figure S2:** Effects of wet *vs* dry surface combined with cooling rate on *CT*_MIN_ and *NT*.

