## Supplementary Material for "A high-throughput method for measuring critical thermal limits of leaves by chlorophyll imaging fluorescence"

**Table S1.** A non-exhaustive sample of the variety of heating or cooling rates for temperature change used for measuring thermal tolerance limits of leaves from the literature.

| **Cold tolerance limits** | |
| --- | --- |
| **Cooling rate** | **Reference** |
| 2 °C h^-1^ | Taschler and Neuner (2004); Neuner and Pramsohler (2006); Sierra-Almeida and Cavieres (2012) |
| 3-4 °C h^-1^ | Robberecht and Junttila (1992) |
| 5 °C h^-1^ | Buchner and Neuner (2009) |
| 10 °C h^-1^ | Menon *et al.* (2015) |
| 1800 °C h^-1^ | Pospı́šil *et al.* (1998) |
| **Heat tolerance limits** | |
| **Heating rate** | **Reference** |
| 30 °C h^-1^ | Frolec *et al.* (2008) |
| 42 °C h^-1^ | Bilger *et al.* (1984) |
| 60 °C h^-1^ | Schreiber and Berry (1977); Schreiber and Armond (1978); (Smillie 1979; Smillie and Nott 1979); Smillie and Gibbons (1981); Bilger *et al.* (1984); Braun *et al.* (2002); Knight and Ackerly (2002); Kim and Portis (2005); Neuner and Pramsohler (2006); Frolec *et al.* (2008); O'Sullivan *et al.* (2013); Tovuu *et al.* (2013); Buchner *et al.* (2015); O'Sullivan *et al.* (2017); Zhu *et al.* (2018) |
| 90 °C h^-1^ | Macias (2011) |
| 120 °C h^-1^ | Bilger *et al.* (1984); Ilík *et al.* (2003); Frolec *et al.* (2008) |
| 180 °C h^-1^ | Frolec *et al.* (2008) |
| 240 °C h^-1^ | Nauš *et al.* (1992) |
| 300 °C h^-1^ | Tovuu *et al.* (2013) |
| 600 °C h^-1^ | Tovuu *et al.* (2013) |
| 648 °C h^-1^ | Tovuu *et al.* (2013) |
| 1800 °C h^-1^ | Pospı́šil *et al.* (1998) |

**Table S2.** Mean ± standard error values for *CT*_MIN_, *CT*_MAX_, and *F*_V_/*F*_M_ for each species and experimental condition (Experiment 2 heating and cooling rates shown for 6, 15, 30, and 60°C h^-1^.

| **Exp. 1**  **Dry surface** | **Species** | | | | | | **Exp. 1**  **Wet surface** | **Species** | | | | | |
| --- | --- | --- | --- | --- | --- | --- | --- | --- | --- | --- | --- | --- | --- |
| **Trait** | ***W. ceracea*** | ***M. citrina*** | | ***E. rubra*** | | ***Q. phellos*** | **Trait** | ***W. ceracea*** | ***M. citrina*** | | ***E. rubra*** | | ***Q. phellos*** |
| *CT*_MIN_ | –13.8 ± 0.4°C | –16.9 ± 0.37°C | | –14.0 ± 0.3°C | | NA | *CT*_MIN_ | –9.8 ± 0.6°C | –13.9 ± 0.4°C | | –9.8 ± 0.8°C | | NA |
| *CT*_MAX_ | 43.2 ± 0.9°C | 46.8 ± 0.8°C | | NA | | 49.2 ± 0.5°C | *CT*_MAX_ | 41.6 ± 0.9°C | 47.2 ± 0.2°C | | NA | | 48.5 ± 0.5°C |
| *F*_V_/*F*_M_ | 0.81 ± 0.004 | 0.77 ± 0.009 | | 0.74 ± 0.008 | | 0.80 ± 0.007 | *F*_V_/*F*_M_ | 0.80 ± 0.004 | 0.78 ± 0.008 | | 0.74 ± 0.006 | | 0.78 ± 0.013 |
| **Exp. 2**  **6°C h^-1^ rate** | **Species** | | | | | | **Exp. 2**  **15°C h^-1^ rate** | **Species** | | | | | |
| **Trait** | ***W. ceracea*** | | ***M. citrina*** | | ***E. rubra*** | | **Trait** | ***W. ceracea*** | | ***M. citrina*** | | ***E. rubra*** | |
| *CT*_MIN_ | –14.1 ± 0.5°C | | –15.2 ± 0.7°C | | –11.7 ± 0.6°C | | *CT*_MIN_ | –14.9 ± 0.2°C | | –14.2 ± 0.5°C | | –11.6 ± 0.5°C | |
| *CT*_MAX_ | 45.8 ± 1.1°C | | 35.3 ± 1.1°C | | 46.7 ± 0.2°C | | *CT*_MAX_ | 42.8 ± 0.5°C | | 38.6 ± 1.0°C | | 44.0 ± 0.3°C | |
| *F*_V_/*F*_M_ | 0.82 ± 0.005 | | 0.73 ± 0.008 | | 0.72 ± 0.005 | | *F*_V_/*F*_M_ | 0.81 ± 0.004 | | 0.74 ± 0.006 | | 0.71 ± 0.006 | |
| **Exp. 2**  **30°C h^-1^ rate** | **Species** | | | | | | **Exp. 2**  **60°C h^-1^ rate** | **Species** | | | | | |
| **Trait** | ***W. ceracea*** | | ***M. citrina*** | | ***E. rubra*** | | **Trait** | ***W. ceracea*** | | ***M. citrina*** | | ***E. rubra*** | |
| *CT*_MIN_ | –12.7 ± 0.6°C | | –12.5 ± 0.6°C | | –12.2 ± 0.4°C | | *CT*_MIN_ | –13.8 ± 0.5°C | | –16.9 ± 0.5°C | | –14.0 ± 0.3°C | |
| *CT*_MAX_ | 41.6 ± 0.4°C | | 41.1 ± 0.5°C | | 44.0 ± 0.4°C | | *CT*_MAX_ | 43.5 ± 0.3°C | | 42.5 ± 0.9°C | | 45.3 ± 0.4°C | |
| *F*_V_/*F*_M_ | 0.83 ± 0.006 | | 0.73 ± 0.006 | | 0.76 ± 0.007 | | *F*_V_/*F*_M_ | 0.82 ± 0.006 | | 0.74 ± 0.011 | | 0.73 ± 0.007 | |

**Table S3.** Full statistical reporting for all species and species-specific effects of wet *vs* dry filter paper surface on *CT*_MIN_ and *CT*_MAX_.

| **Response: *CT*_MIN_** | **All species** | | | ***W. ceracea*** | | | ***M. citrina*** | | | ***E. rubra*** | | |
| --- | --- | --- | --- | --- | --- | --- | --- | --- | --- | --- | --- | --- |
| **Fixed effects** | **Estimate** | **95% CI** | ***p*** | **Estimate** | **95% CI** | ***p*** | **Estimate** | **95% CI** | ***p*** | **Estimate** | **95% CI** | ***p*** |
| Dry surface /  *E. rubra* (intercept) | –18.36 | –32.49 - –4.22 | **0.011** | Intercept:  –5.71 | –50.92 - 39.49 | 0.800 | Intercept:  –20.36 | –35.80 - –4.93 | **0.012** | Intercept:  –31.26 | –54.11 - –8.41 | **0.009** |
| Wet surface | 3.81 | 2.77 - 4.85 | **<0.001** | 3.92 | 1.99 - 5.86 | **<0.001** | 2.98 | 1.31 - 4.65 | **0.001** | 3.99 | 2.34 - 5.63 | **<0.001** |
| *F*_V_/*F*_M_ | 6.19 | –12.89 - 25.27 | 0.521 | –9.89 | –64.94 - 45.15 | 0.719 | 4.72 | –16.14 - 25.58 | 0.645 | 23.54 | –7.53 - 54.60 | 0.132 |
| *M. citrina* | –3.50 | –4.92 - –2.07 | **<0.001** | -- | -- | -- | -- | -- | -- | -- | -- | -- |
| *W. ceracea* | –0.42 | –2.36 - 1.51 | 0.664 | -- | -- | -- | -- | -- | -- | -- | -- | -- |
| R^2^ | 0.464 | -- | -- | 0.288 | -- | -- | 0.374 | -- | -- | 0.527 | -- | -- |
| **Response: *CT*_MAX_** | **All species** | | | ***W. ceracea*** | | | ***M. citrina*** | | | ***Q. phellos*** | | |
| **Fixed effects** | **Estimate** | **95% CI** | ***p*** | **Estimate** | **95% CI** | ***p*** | **Estimate** | **95% CI** | ***p*** | **Estimate** | **95% CI** | ***p*** |
| Dry surface /  *M. citrina* (intercept) | 32.76 | 18.03 - 47.49 | **<0.001** | Intercept:  6.90 | –27.59 - 41.40 | 0.683 | Intercept:  36.31 | 4.91 - 67.71 | **0.025** | Intercept:  47.34 | 31.89 - 62.79 | **<0.001** |
| Wet surface | –0.55 | –1.63 - 0.54 | 0.317 | –1.47 | –3.93 - 1.00 | 0.232 | 0.32 | –1.33 - 1.97 | 0.694 | –0.63 | –2.08 - 0.82 | 0.379 |
| *F*_V_/*F*_M_ | 18.20 | –0.12 - 36.52 | 0.052 | 46.02 | 2.33 - 89.71 | **0.040** | 13.16 | –26.19 - 52.50 | 0.497 | 2.32 | –17.03 - 21.68 | 0.806 |
| *Q. phellos* | 2.01 | 0.68 - 3.35 | **0.004** | -- | -- | -- | -- | -- | -- | -- | -- | -- |
| *W. ceracea* | –4.47 | –5.80 - –3.14 | **<0.001** | -- | -- | -- | -- | -- | -- | -- | -- | -- |
| R^2^ | 0.593 | -- | -- | 0.213 | -- | -- | 0.028 | -- | -- | 0.041 | -- | -- |

**Table S4.** Full statistical reporting for effects of wet *vs* dry surface in combination with cooling rate on *CT*_MIN_.

| **Response: *CT*_MIN_** | **All species** | | | ***W. ceracea*** | | | ***M. citrina*** | | | ***E. rubra*** | | |
| --- | --- | --- | --- | --- | --- | --- | --- | --- | --- | --- | --- | --- |
| **Fixed effects** | **Estimate** | **95% CI** | ***p*** | **Estimate** | **95% CI** | ***p*** | **Estimate** | **95% CI** | ***p*** | **Estimate** | **95% CI** | ***p*** |
| Dry surface /  Rate = 15 °C h^-1^ /  *E. rubra* (Intercept) | –11.49 | –21.77 - –2.01 | **0.019** | Intercept:  –4.23 | –43.83 - 35.37 | 0.832 | Intercept:  –12.39 | –24.09 - –0.69 | **0.038** | Intercept:  –20.60 | –32.47 - –8.74 | **0.001** |
| Wet surface | 4.72 | 3.45 - 5.99 | **<0.001** | 6.42 | 3.90 - 8.94 | **<0.001** | 4.03 | 2.10 - 5.95 | **<0.001** | 3.73 | 2.17 - 5.30 | **<0.001** |
| Rate = 60 °C h^-1^ | –1.11 | –2.14 - –0.08 | **0.034** | 1.17 | –0.77 - 3.11 | 0.235 | –2.69 | –4.13 - –1.25 | **<0.001** | –2.70 | –4.15 - –1.26 | **<0.001** |
| Wet surface ×  rate = 60 °C h^-1^ | –0.90 | –2.53 - 0.73 | 0.279 | –2.53 | –5.74 - 0.68 | 0.121 | –0.98 | –3.42 - 1.45 | 0.420 | 0.34 | –1.81 - 2.48 | 0.754 |
| *F*_V_/*F*_M_ | –0.55 | –14.28 - 13.19 | 0.938 | –13.12 | –61.73 - 35.48 | 0.592 | –2.44 | –18.29 - 13.41 | 0.758 | 12.71 | –3.99 - 29.40 | 0.133 |
| *M. citrina* | –3.07 | –4.11 - –2.02 | **<0.001** | -- | -- | -- | -- | -- | -- | -- | -- | -- |
| *W. ceracea* | –0.95 | –2.50 - 0.59 | 0.225 | -- | -- | -- | -- | -- | -- | -- | -- | -- |
| Marginal R^2^ | 0.458 | -- | -- | 0.387 | -- | -- | 0.538 | -- | -- | 0.552 | -- | -- |

**Table S5.** Full statistical reporting for all species and species-specific effects of variable cooling rate on *CT*_MIN_.

| **Response: *CT*_MIN_** | **All species** | | | ***W. ceracea*** | | | ***M. citrina*** | | | ***E. rubra*** | | |
| --- | --- | --- | --- | --- | --- | --- | --- | --- | --- | --- | --- | --- |
| **Fixed effects** | **Estimate** | **95% CI** | ***p*** | **Estimate** | **95% CI** | ***p*** | **Estimate** | **95% CI** | ***p*** | **Estimate** | **95% CI** | ***p*** |
| Rate = 3 °C h^-1^ /  *E. rubra* (Intercept) | –11.38 | –17.01 - –5.75 | **<0.001** | –40.89 | –70.67 - –11.10 | **0.008** | –16.82 | –25.42 - –8.21 | **<0.001** | –11.58 | –19.21 - –3.96 | **0.003** |
| Rate = 6 °C h^-1^ | –0.33 | –1.14 - 0.48 | 0.422 | 0.62 | –0.67 - 1.92 | 0.343 | –1.81 | –3.49 - –0.12 | **0.036** | –0.15 | –1.29 - 0.98 | 0.790 |
| Rate = 15 °C h^-1^ | –0.32 | –1.12 - 0.49 | 0.438 | –0.12 | –1.42 - 1.19 | 0.859 | –0.80 | –2.43 - 0.84 | 0.335 | –0.10 | –1.22 - 1.03 | 0.864 |
| Rate = 30 °C h^-1^ | 0.75 | –0.05 - 1.55 | 0.065 | 1.67 | 0.37 - 2.97 | **0.012** | 0.91 | –0.70 - 2.51 | 0.265 | –0.74 | –1.93 - 0.45 | 0.220 |
| Rate = 60 °C h^-1^ | –1.34 | –2.15 - –0.53 | **0.001** | 0.75 | –0.47 - 1.97 | 0.225 | –3.51 | –5.19 - –1.82 | **<0.001** | –2.47 | –3.69 - –1.25 | **<0.001** |
| Rate = 240 °C h^-1^ | –0.80 | –1.60 - 0.01 | 0.052 | 0.70 | –0.62 - 2.02 | 0.298 | –1.74 | –3.35 - –0.13 | **0.035** | –1.90 | –3.11 - –0.70 | **0.002** |
| *F*_V_/*F*_M_ | –0.89 | –8.55 - 6.78 | 0.82 | 32.04 | –4.23 - 68.32 | 0.083 | 4.66 | –6.84 - 16.16 | 0.422 | 0.12 | –10.42 - 10.66 | 0.982 |
| *M. citrina* | –2.18 | –2.76 - –1.60 | **<0.001** | -- | -- | -- | -- | -- | -- | -- | -- | -- |
| *W. ceracea* | –1.50 | –2.36 - –0.63 | **0.001** | -- | -- | -- | -- | -- | -- | -- | -- | -- |
| Marginal R^2^ | 0.230 | -- | -- | 0.126 | -- | -- | 0.332 | -- | -- | 0.220 | -- | -- |

**Table S6.** Full statistical reporting for all species and species-specific effects of variable heating rate on *CT*_MAX_.

| **Response: *CT*_MAX_** | **All species** | | | ***W. ceracea*** | | | ***M. citrina*** | | | ***E. rubra*** | | |
| --- | --- | --- | --- | --- | --- | --- | --- | --- | --- | --- | --- | --- |
| **Fixed effects** | **Estimate** | **95% CI** | ***p*** | **Estimate** | **95% CI** | ***p*** | **Estimate** | **95% CI** | ***p*** | **Estimate** | **95% CI** | ***p*** |
| Rate = 60 °C h^-1^ /  *E. rubra* (Intercept) | 27.79 | 20.09 - 35.50 | **<0.001** | Intercept:  14.87 | –0.48 - 30.22 | 0.058 | Intercept:  27.79 | 18.26 - 37.32 | **<0.001** | Intercept:  41.75 | 31.47 - 52.03 | **<0.001** |
| Rate = 6 °C h^-1^ | –0.68 | –1.86 - 0.50 | 0.259 | 1.60 | 0.24 - 2.96 | **0.021** | –7.71 | –9.48 - –5.93 | **<0.001** | 1.38 | –0.19 - 2.94 | 0.085 |
| Rate = 15 °C h^-1^ | –2.43 | –3.62 - –1.24 | **<0.001** | –2.00 | –3.47 - –0.52 | **0.008** | –4.68 | –6.43 - –2.94 | **<0.001** | –1.40 | –2.98 - 0.17 | 0.080 |
| Rate = 30 °C h^-1^ | –1.74 | –2.72 - –0.77 | **0.001** | –2.11 | –3.18 - –1.03 | **<0.001** | –2.10 | –3.66 - –0.54 | **0.009** | –1.31 | –2.61 - –0.01 | **0.048** |
| Rate = 45 °C h^-1^ | –1.48 | –2.44 - –0.51 | **0.003** | –0.72 | –1.80 - 0.36 | 0.188 | –0.95 | –2.53 - 0.62 | 0.232 | –2.68 | –3.96 - –1.40 | **<0.001** |
| Rate = 120 °C h^-1^ | 1.00 | –0.06 - 2.07 | 0.065 | 1.78 | 0.62 - 2.95 | **0.003** | 2.24 | 0.48 - 3.99 | **0.013** | –0.45 | –1.95 - 1.05 | 0.554 |
| Rate = 240 °C h^-1^ | 2.03 | 0.95 - 3.12 | **<0.001** | 2.78 | 1.58 - 3.98 | **<0.001** | 3.76 | 1.86 - 5.67 | **<0.001** | –0.13 | –1.56 - 1.29 | 0.853 |
| *F*_V_/*F*_M_ | 23.79 | 12.68 - 34.91 | **<0.001** | 38.48 | 17.91 - 59.05 | **<0.001** | 21.98 | 7.80 - 36.16 | **0.003** | 5.15 | –9.68 - 19.99 | 0.492 |
| *M. citrina* | –1.30 | –1.96 - –0.63 | **<0.001** | -- | -- | -- | -- | -- | -- | -- | -- | -- |
| *W. ceracea* | –1.24 | –2.02 - –0.45 | **0.002** | -- | -- | -- | -- | -- | -- | -- | -- | -- |
| Marginal R^2^ | 0.429 | -- | -- | 0.619 | -- | -- | 0.863 | -- | -- | 0.319 | -- | -- |

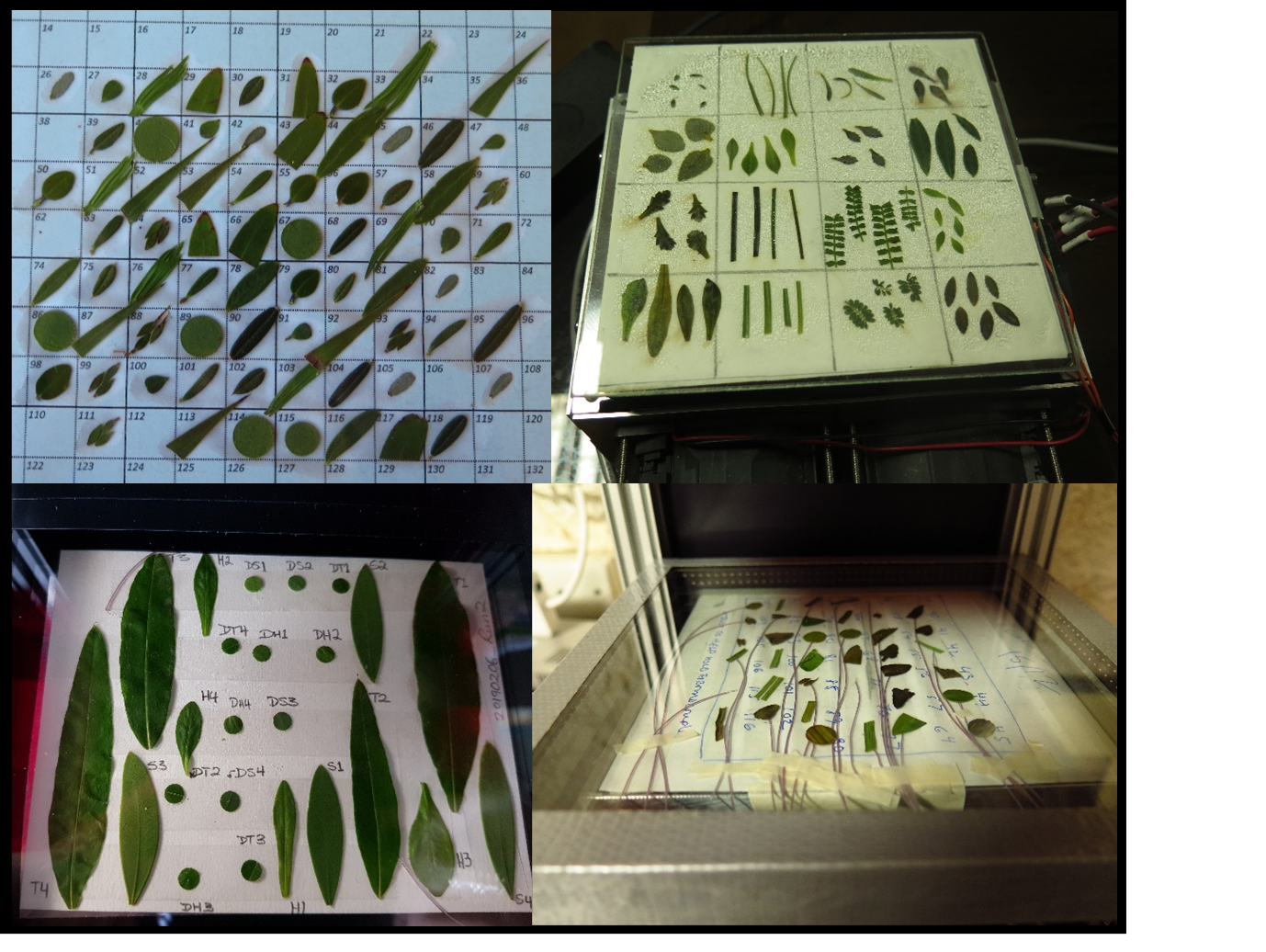

**Fig. S1.** Various experimental applications of the Peltier plate and chlorophyll fluorescence Maxi-Imaging-PAM system using whole leaves, leaf sections, and leaf discs of multiple species, and the potential application of type-T thermocouples for recording the temperature of individual samples. Images taken by Verónica F. Briceño and Pieter A. Arnold.

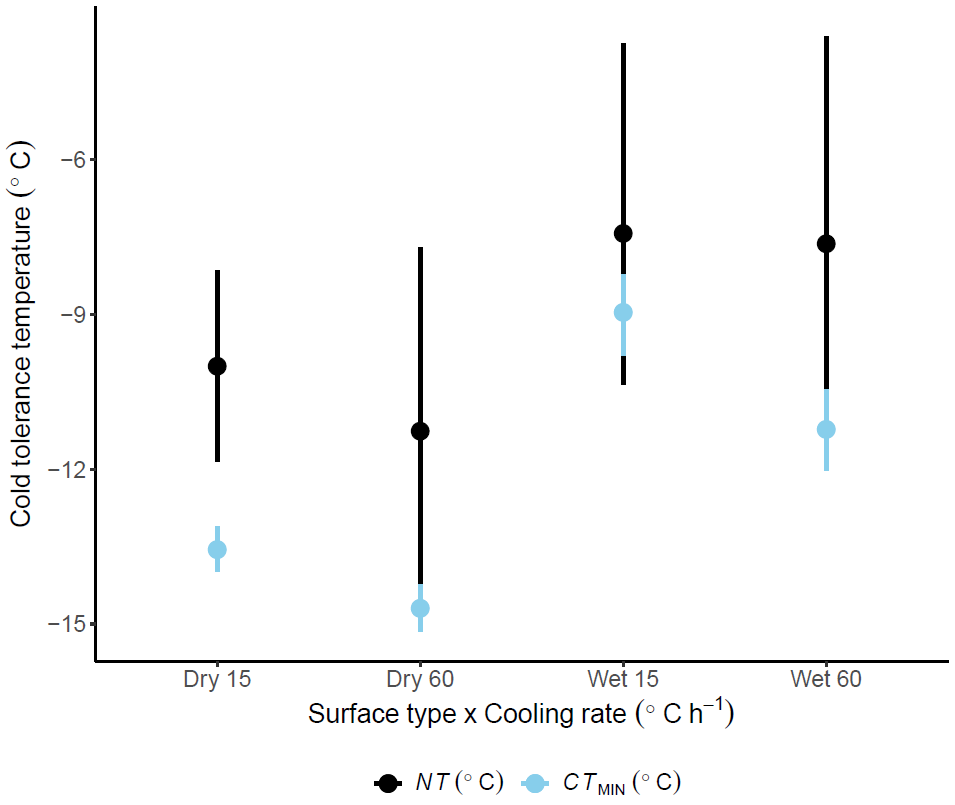

**Fig. S2.** The effect of varying cooling rate (°C h^-1^) in combination with varying surfaces (dry *vs* wet filter paper) on the *NT* (black circles) and *CT*_MIN_ (light blue circles) estimates (°C). *NT* was measured only on a small, random subset of leaves using the two thermocouples attached to leaves from various species on the Peltier plate (total *n* = 17, therefore the 95% CIs are much larger than those of *CT*_MIN_. For consistency in comparison, the *CT*_MIN_ values are also grouped across the three species.
